# Tobacco Carcinogen Induces Persistent DNA Damage and Homologous Recombination Dysregulation in Pancreatic Ductal Carcinogenesis

**DOI:** 10.64898/2026.10.13.758888

**Authors:** Varunkumar Krishnamoorthy, Vineet K. Gupta, Sudhakar Jinka, Supriya Srinivasan, Kumaraswamy Honnenahally, Chanjuan Shi, Poorva Poorva, X. Steven Chen, Nipun B. Merchant, Austin R. Dosch, Nagaraj S. Nagathihalli

## Abstract

Cigarette smoking is a well-established risk factor for the development of pancreatic ductal adenocarcinoma (PDAC), yet the mechanisms by which tobacco carcinogens, such as Nicotine-Derived Nitrosamine Ketone (NNK), drive pancreatic carcinogenesis remain incompletely understood. We investigated the impact of NNK on pancreatic epithelium by treating human pancreatic ductal epithelial cells (HPDE6-E6E7; H6c7) with NNK for 5 or 50 days followed by genome-wide RNA sequencing, qRT-PCR, immunofluorescence, comet assay, and γH2AX ELISA. We assessed γH2AX by immunohistochemistry in human PDAC tissue microarrays and in TCGA-PAAD datasets to identify clinical correlates. Acute NNK exposure (5 days) induced DNA double-strand breaks and activated ATM-γH2AX signaling, accompanied by increased RAD51 and BRCA2 recruitment, consistent with activation of homologous recombination (HR)-mediated DNA repair. In contrast, chronic NNK exposure (50 days) resulted in persistent DNA damage signaling and transcriptional upregulation of HR-associated genes, including RAD51, BRCA1, BRCA2, FANCD2, and PLK1. Despite sustained accumulation of RAD51- and BRCA2-positive repair foci, chronic NNK exposure was associated with altered RAD51 localization, persistent DNA damage, and chromosomal instability, suggesting dysregulation of HR repair fidelity during prolonged genotoxic stress. γH2AX expression was significantly elevated in NNK-treated cells, KRAS-driven premalignant pancreatic lesions, and pancreatic tissues from smokers. Increased γH2AX expression correlated with tumor progression and poor overall survival, while elevated RAD51 expression was associated with adverse outcomes in patients with PDAC. Collectively, these findings reveal a biphasic DNA damage response to NNK characterized by early activation of ATM-γH2AX-dependent DNA repair followed by persistent DNA damage signaling, dysregulated HR repair, and genomic instability during chronic exposure. Tobacco-induced disruption of DNA repair homeostasis may represent a key mechanism driving pancreatic carcinogenesis and supports γH2AX as a clinically relevant prognostic biomarker in tobacco-associated PDAC.

## INTRODUCTION

Pancreatic ductal adenocarcinoma (PDAC) remains one of the deadliest malignancies and is projected to become one of the leading causes of cancer-related deaths worldwide. Despite advances in surgery, chemotherapy, and targeted interventions, the overall five-year survival rate remains dismal due to aggressive disease biology, late-stage diagnosis, and profound therapeutic resistance (1, 2). Consequently, identifying modifiable risk factors and understanding the molecular mechanisms that drive pancreatic tumor initiation and progression remain critical priorities.

Cigarette smoking is among the most established environmental risk factors for PDAC and is estimated to account for approximately 20-25% of pancreatic cancer cases (3, 4). Numerous epidemiological studies have demonstrated that current smokers exhibit a significantly elevated risk of developing pancreatic cancer compared with never-smokers, while smoking cessation gradually reduces this risk over time (4, 5). Although the carcinogenic effects of tobacco exposure are strongly associated with pancreatic cancer incidence, the molecular pathways through which tobacco-derived carcinogens promote pancreatic tumorigenesis remain incompletely understood.

More than 7,000 chemicals are present in cigarette smoke; 4-(Methylnitrosamino)-1-(3-pyridyl)-1-butanone (Nicotine-Derived Nitrosamine Ketone or NNK) is one of the most potent tobacco-specific nitrosamines and a well-characterized carcinogen implicated in smoking-associated malignancies, including lung and pancreatic cancers. NNK undergoes metabolic activation to generate DNA-reactive intermediates that can induce DNA adduct formation, oxidative stress, genomic instability, and oncogenic mutations (6, 7). Importantly, NNK has been shown to accelerate Kras-driven pancreatic neoplasia in genetically engineered mouse models, providing direct evidence linking tobacco exposure to pancreatic carcinogenesis (8, 9).

Genomic instability is a hallmark of cancer and frequently arises from persistent DNA damage and defective DNA repair responses (10). The DNA damage response (DDR) network is a critical safeguard against genomic instability by sensing DNA lesions, coordinating cell-cycle checkpoints, and activating DNA repair pathways (10). Among the earliest events following DNA double-strand break (DSB) formation is phosphorylation of histone H2AX at serine-139 (γH2AX), a sensitive marker of DNA damage that facilitates recruitment of repair complexes (11–13). Activation of the ATM kinase further amplifies DDR signaling by phosphorylating multiple downstream effectors involved in DNA repair, checkpoint regulation, and cell survival (14). Dysregulated DDR signaling has been increasingly implicated in pancreatic cancer development, progression, and therapeutic resistance (15, 16).

Homologous recombination (HR) represents a high-fidelity DSB repair mechanism orchestrated by key mediators including BRCA1, BRCA2, RAD51, PALB2, and FANCD2 (17–19). Defects in HR repair pathways are common in PDAC and have emerged as clinically important biomarkers that predict sensitivity to platinum-based therapies and PARP inhibitors (20–22). However, while inherited and somatic alterations in DDR genes have been extensively characterized in pancreatic cancer, relatively little is known regarding how tobacco carcinogens influence DDR activation and HR repair during the earliest stages of pancreatic tumorigenesis.

Emerging evidence suggests that chronic carcinogen exposure can induce persistent DNA damage and adaptive activation of DNA repair pathways, thereby enabling damaged cells to survive and accumulate oncogenic mutations (10, 23). Whether prolonged NNK exposure promotes sustained DDR signaling, enhances homologous recombination repair activity, and contributes to pancreatic neoplastic progression remains unclear. Moreover, the clinical significance of tobacco-associated DDR activation in pancreatic cancer patients has not been fully elucidated.

In the present study, we investigated the impact of acute and chronic NNK exposure on DNA damage signaling and homologous recombination repair in pancreatic epithelial cells, PanIN (pancreatic intraepithelial neoplasia) cells, Kras-driven pancreatic cancer mouse models, and human pancreatic tissues. Our findings show that chronic NNK exposure induces persistent activation of the ATM/γH2AX axis and increases expression of key HR repair proteins, including RAD51 and BRCA2, thereby promoting pancreatic neoplastic progression. Furthermore, elevated DDR activation observed in pancreatic tumors from smokers and its association with adverse clinical outcomes suggests that aberrant DNA damage signaling represents a critical mechanistic link between tobacco exposure and pancreatic tumorigenesis. Collectively, these results provide new insights into smoking-associated pancreatic cancer pathogenesis and identify DDR-associated pathways as potential therapeutic targets in tobacco-driven PDAC.

## RESULTS

### Chronic NNK exposure drives a compensatory transcriptional upregulation of homologous recombination repair genes

In pancreatic cancer, cigarette smoking is a major risk factor. Our previously published data confirm that smokers have lower survival when compared to non-smokers in human patient datasets (8). To test whether NNK-induced DNA damage elicits a transcriptional HR response, we performed genome-wide RNA-seq of H6c7 cells treated with NNK (1 µmol/L) for 5 days (acute) or 50 days (chronic). Cuffdiff analysis (q < 0.05) identified 1,077 differentially expressed genes at 5 days and 6,166 at 50 days, reflecting a markedly expanded transcriptional response with prolonged NNK exposure (Fig. 1A). Genome-wide ranked waterfall and distribution plots further illustrate the magnitude and distribution of these NNK-induced transcriptional alterations (Supplementary Fig. 1A, B). None of the core HR genes RAD51, BRCA1, BRCA2, FANCD2, or PLK1 were significantly altered at 5 days. In contrast, chronic 50-day NNK exposure induced robust upregulation of the entire HR machinery: RAD51 (3.36-fold), BRCA1 (3.06-fold), PLK1 (2.98-fold), BRCA2 (2.79-fold), FANCD2 (1.97-fold), XRCC1 (2.03-fold), PARP1 (1.69-fold), PCNA (1.62-fold), PALB2 (1.49-fold), and CHEK2 (1.42-fold) (all q < 0.05; Fig. 1C). These findings were corroborated by qRT-PCR of H6c7 cells treated with NNK for 50 days, demonstrating significant upregulation of BRCA1, BRCA2, FANCD2, PLK1, and RAD51 mRNA (all p <0.01; Fig. 1D). The RNA-seq dataset identified DNA damage response, homologous recombination, and cell cycle checkpoint signaling as the top enriched pathways (PANTHER and Ingenuity analyses; see Methods), with Fig. 1E providing a schematic overview of this NNK-engaged DNA damage response pathway.

**Figure 1.**
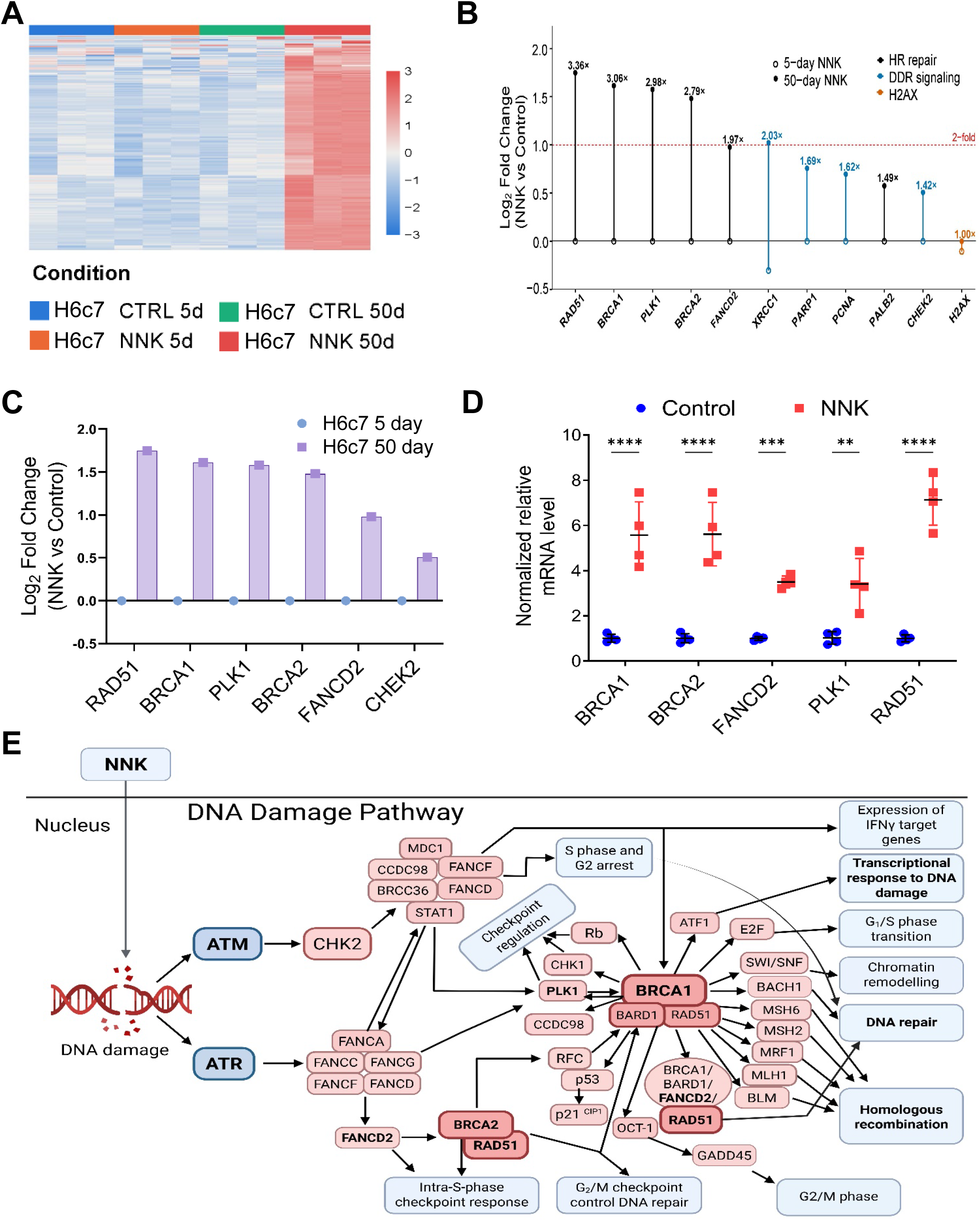
NNK induces a two-phase DNA damage response in pancreatic epithelial cells. (**A**) Heatmap of row-scaled top 200 gene expression (blue-white-red scale) across four H6c7 cell conditions: 5-day control, 5-day NNK, 50-day control, and 50-day NNK. (**B**) Temporal progression plot showing log₂ fold change of 11 DDR genes in H6c7 cells at 5-day (open circles) and 50-day (filled circles) NNK treatment. Lines connect time points for each gene. Genes are color-coded by functional class: HR repair (black; RAD51, BRCA1, PLK1, BRCA2, FANCD2), DDR signaling (teal; XRCC1, PARP1, PCNA, PALB2, CHEK2), and H2AX (orange). Fold-change values are annotated for significant 50-day changes. (**C**) Bar chart of log₂ fold change (NNK vs. control) for six DDR genes in H6c7 cells at 5-day (circles) and 50-day (bars) NNK treatment. H2AX is unchanged at either time point (see **B**). (**D**) qRT-PCR validation of BRCA1, BRCA2, FANCD2, PLK1, and RAD51 mRNA expression in H6c7 cells following 50-day NNK treatment. Individual data points are shown with mean ± SEM; significance was determined by Student’s t-test. (**E**) Schematic of the DNA damage response pathway engaged by NNK, highlighting ATM/ATR signaling and downstream homologous recombination effectors (created with BioRender). (**p < 0.01; ***p < 0.001; ****p < 0.0001).

### NNK induces DNA double-strand breaks and activates the DNA damage response in human pancreatic ductal epithelial cells

To determine whether NNK directly induces genomic damage, we exposed pancreatic ductal epithelial cells to NNK for 5 days and evaluated DNA damage. Comet assay analysis revealed a dose-dependent increase in DNA strand breaks in H6c7 cells following NNK treatment, indicating rapid induction of genotoxic stress (Fig. 2A). Consistent with these findings, ELISA-based quantification demonstrated a concentration-dependent increase in γH2AX levels following NNK exposure, confirming the accumulation of DNA double-strand breaks in response to acute carcinogen treatment (Fig. 2B). To validate DNA damage at the cellular level, γH2AX immunofluorescence (IF) staining was performed in H6c7 and PanIN cells after 5 days of NNK exposure. Compared with controls, NNK-treated cells exhibited a marked increase in nuclear γH2AX-positive foci, demonstrating activation of the DNA damage response in both normal (H6c7) and premalignant (PanIN) pancreatic epithelial cells (Fig. 2C). We next examined whether chronic NNK exposure results in sustained DNA damage signaling. Following 50 days of NNK treatment, IF analysis revealed an increase in both pATM and γH2AX-positive nuclei compared with control cells (Fig. 2D). Co-localization of pATM and γH2AX signals further indicated the activation of the ATM-dependent DNA damage response pathway during chronic carcinogen exposure. We further examined somatic mutation frequencies of DDR genes across four publicly available PDAC cohorts; ATM was the most frequently mutated DDR gene (Supplementary Fig. 2A&B). Collectively, these findings demonstrate that NNK rapidly induces DNA double-strand breaks and activates the DNA damage response signaling in pancreatic epithelial cells, while chronic exposure sustains ATM-mediated checkpoint signaling, supporting a role for chronic NNK exposure in promoting genomic stress.

**Figure 2.**
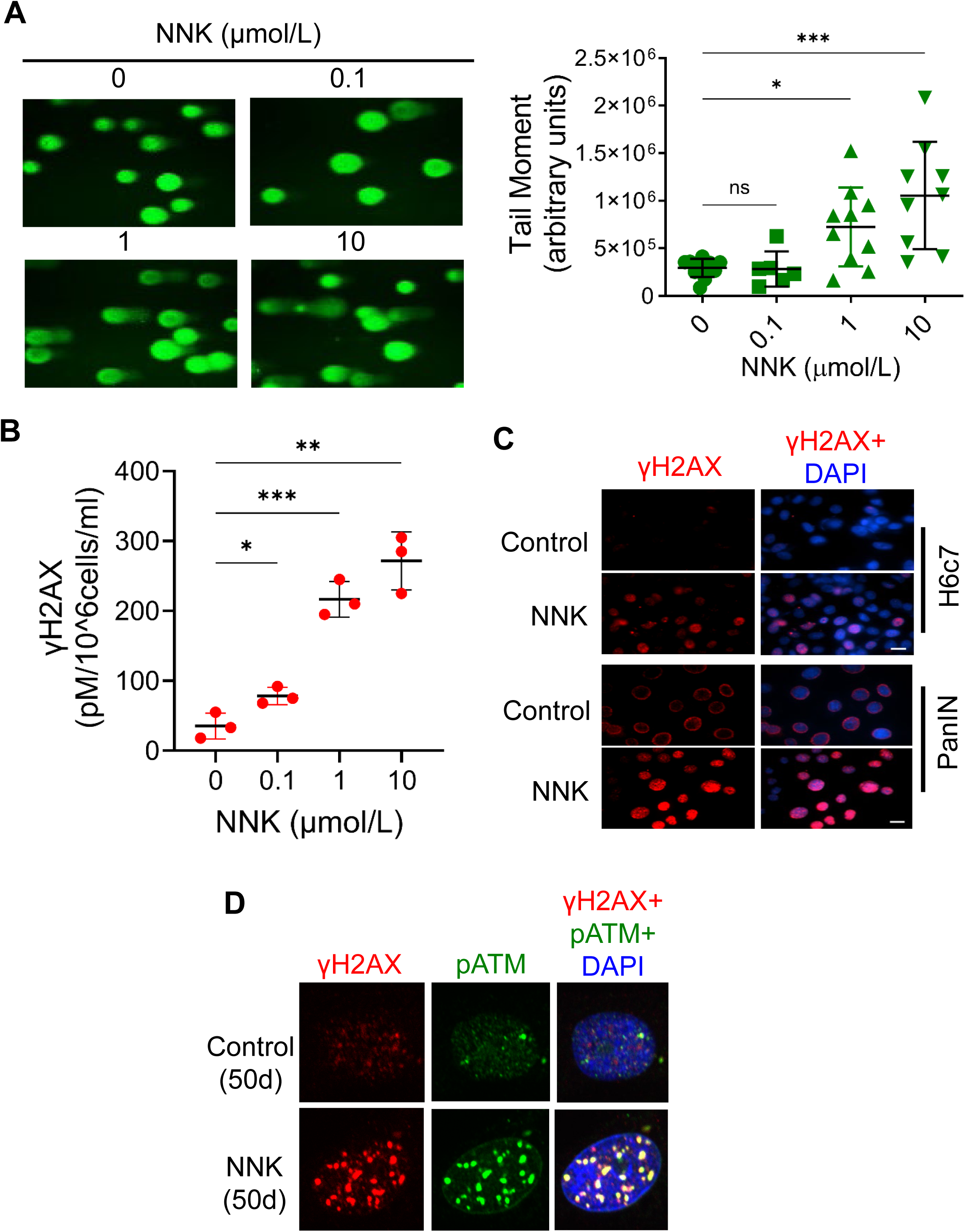
NNK induces DNA double-strand breaks and activates ATM–γH2AX signaling in pancreatic cells. **A**) Neutral comet assay in H6c7 cells treated with NNK at 0, 0.1, 1, and 10 μmol/L. Representative images and tail moment quantification are shown. Data are the mean ± SEM from three independent experiments; significance was determined by one-way ANOVA with Tukey’s post-hoc test. (**B**) γH2AX ELISA in H6c7 cells following dose-dependent NNK treatment (0–10 μmol/L). Data are mean ± SEM; significance determined by one-way ANOVA. (**C**) Immunofluorescence imaging of γH2AX foci in H6c7 and PanIN cells following NNK treatment versus DMSO control. Representative images are shown; scale bar, 10 μm. (**D**) Co-immunofluorescence of pATM (Ser1981) and γH2AX in H6c7 cells following 50-day NNK treatment, demonstrating co-localization at DSB sites. Data are mean ± SEM; significance determined by Student’s *t*-test. (*p < 0.05; **p < 0.01; ***p < 0.001; ns, not significant).

### Tobacco smoking is associated with elevated γH2AX expression in KC mice and human pancreatic tissues

To investigate whether the tobacco carcinogens accelerate pancreatic neoplasia in vivo, *Ptf1a^CreERTM/+^;LSL-Kras^G12D/+^*(*KC*) mice were established as a genetically engineered premalignant model of pancreatic cancer. Unlike more aggressive PDAC models such as KPC mice, KC mice develop acinar-to-ductal metaplasia and pancreatic intraepithelial neoplasia (PanIN) lesions with limited progression to invasive carcinoma, thereby closely recapitulating the earliest stages of human pancreatic tumorigenesis. This extended premalignant phase provides a unique opportunity to investigate mechanisms that promote pancreatic cancer initiation and progression, making KC mice particularly suitable for studying environmental carcinogens, DNA damage responses, and cancer prevention strategies. We chronically exposed KC mice to NNK (Fig. 3A), and pancreatic tissues were evaluated histologically. NNK-treated KC mice displayed enhanced PanIN development accompanied by increased Alcian blue staining, elevated CK19 expression, and marked accumulation of γH2AX-positive lesions relative to controls. These observations demonstrate that chronic NNK exposure promotes neoplastic progression in Kras-driven pancreatic tumorigenesis while maintaining a persistent DNA damage state (Fig. 3B, C). To examine the relationship between smoking and DNA damage signaling in human pancreatic tissues, γH2AX was evaluated by immunohistochemistry in a tissue microarray containing normal duct, chronic pancreatitis, and tumor specimens. Among evaluable cores with smoking status and weak/strong γH2AX classification, 76 were from smokers and 50 from non-smokers (Fig. 3D, E). Strong γH2AX staining was more frequent among smokers across the three tissue categories. Using the counts displayed in Fig. 3E, a Cochran-Mantel-Haenszel analysis stratified by tissue type yielded a common odds ratio of 7.85 for strong versus weak staining in smokers compared with non-smokers (95% CI, 3.34-18.45; P < 0.000001). These human tissue data are consistent with greater γH2AX-associated DNA damage signaling among smokers, while the mouse data support persistent DDR activation during NNK-accelerated PanIN progression.

**Figure 3.**
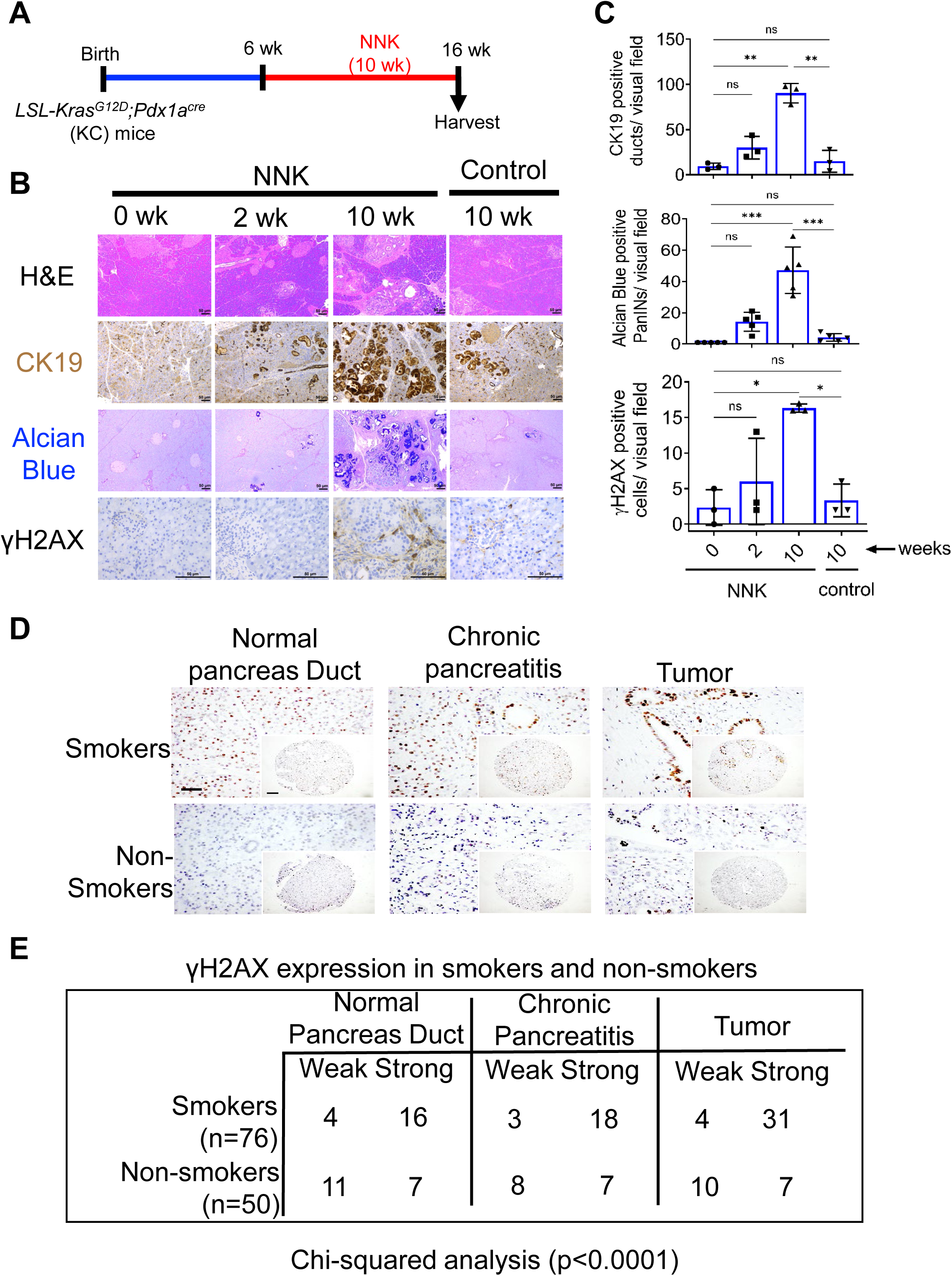
Tobacco smoking is associated with elevated γH2AX in mouse and human pancreatic tissues. **(A)** Schematic representation of treatment time points in KC (LSL-Kras^G12D/+^; Pdx1a^Cre/+^) mice. **(B)** Representative hematoxylin and eosin (H&E), CK19, Alcian Blue, and γH2AX immunohistochemical staining of pancreatic tissue from KC mice treated with NNK at 0, 2, and 10 weeks. PanIN lesion progression is shown. Scale bar, 50 μm. (**C**) Quantification of CK19-positive ducts, Alcian Blue–positive PanIN lesions, and γH2AX-positive cells per visual field in KC mice at 0, 2, and 10 weeks of NNK treatment versus 10-week controls. Data are mean ± SEM. (**D**) Representative immunohistochemical images of γH2AX staining in normal pancreatic duct, chronic pancreatitis, and tumor tissue from smokers and non-smokers. Tissue microarray (TMA; n = 136 cores). Scale bar, 50 μm. (E) Contingency table of γH2AX staining (weak vs. strong) in normal duct, chronic pancreatitis, and tumor cores from smokers (n = 76) versus non-smokers (n = 50). Significance determined by chi-square test (*p < 0.05; **p < 0.01; ***p < 0.001; ns, not significant).

### NNK impairs homologous recombination fidelity and induces chromosomal instability

After establishing that NNK induces DNA damage, we next investigated whether NNK activates homologous recombination (HR) repair pathways. Quantification of γH2AX foci revealed a significant increase in DNA damage following NNK treatment compared with control cells (Fig. 4A). Pharmacologic inhibition of ATM using KU55933 markedly reduced NNK-induced γH2AX accumulation, indicating that ATM activity contributes to the DNA damage response provoked by acute NNK exposure. To examine the kinetics of DNA damage, we monitored γH2AX-positive cells over time following NNK exposure. NNK-treated cells exhibited a sustained increase in γH2AX positivity relative to controls, whereas ATM inhibition significantly attenuated the accumulation and persistence of γH2AX-positive cells throughout the time course (Fig. 4B). These findings indicate that ATM signaling is a critical mediator of the NNK-induced DNA damage response. Because RAD51 is a central effector of HR-mediated DNA repair, we next assessed its subcellular localization following NNK treatment. Cellular fractionation and immunoblot analyses showed a dose-dependent increase in nuclear RAD51 accumulation with increasing NNK concentrations, accompanied by a corresponding reduction in cytoplasmic RAD51 levels (Fig. 4C), indicating NNK-induced RAD51 nuclear translocation. Consistent with these observations, immunofluorescence analysis revealed enhanced nuclear co-localization of RAD51 and BRCA2 in NNK-treated cells compared with controls (Fig. 4D). Increased RAD51-BRCA2 foci formation indicates activation of the HR repair machinery in response to NNK-induced DNA damage. To test whether ATM regulates RAD51 recruitment, we quantified RAD51 foci formation in the presence or absence of KU55933. NNK treatment significantly increased RAD51 foci formation, whereas ATM inhibition markedly reduced RAD51 accumulation despite NNK exposure (Fig. 4E). These findings show that NNK-induced DNA damage activates the HR pathway by increasing nuclear accumulation of RAD51 and BRCA2, supporting a compensatory DNA repair response to tobacco carcinogen-induced genomic stress.

**Figure 4.**
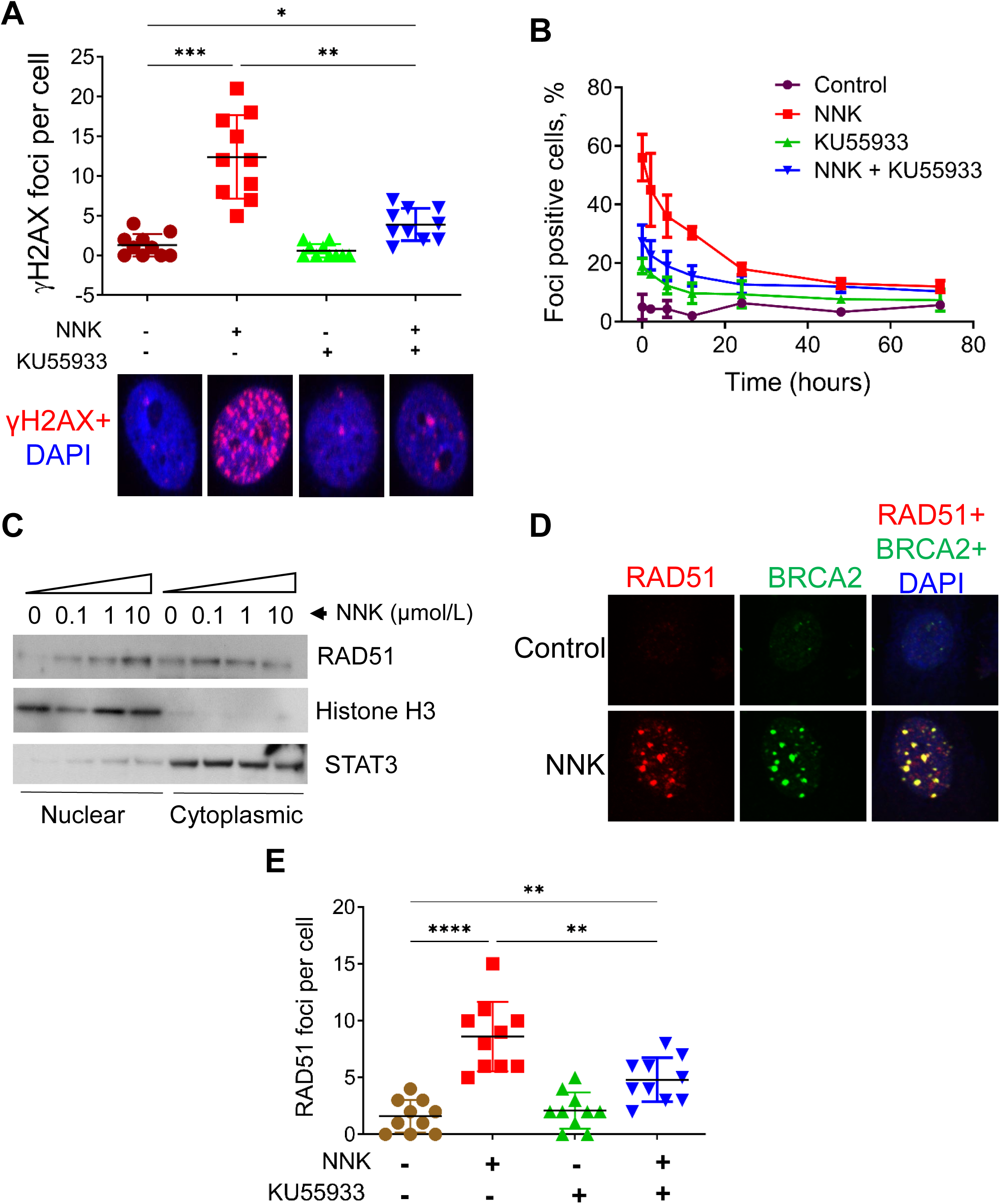
Chronic NNK impairs ATM-dependent DSB repair and induces RAD51 mislocalization. **(A**) Quantification of γH2AX foci per nucleus in H6c7 cells following NNK treatment, with and without the ATM inhibitor KU55933 (10 μmol/L). Data are mean ± SEM from three independent experiments; significance was determined by two-way ANOVA. (**B**) Time course of γH2AX foci–positive cells (%) over 72 hours following NNK treatment, with or without KU55933 (10 μmol/L). Data are mean ± SEM. (**C**) Subcellular fractionation western blot showing nuclear and cytoplasmic RAD51, Histone H3, and STAT3 protein levels in H6c7 cells following increasing NNK concentrations (0–10 μmol/L). (**D**) Co-immunofluorescence of BRCA2 and RAD51 in H6c7 cells following 50-day NNK treatment, demonstrating enhanced nuclear co-localization of RAD51 and BRCA2. Data are mean ± SEM; significance determined by Student’s *t*-test (*p < 0.05; **p < 0.01; ***p < 0.001; ****p < 0.0001). (**E**) Quantification of RAD51 foci per cell in H6c7 cells treated with NNK, with or without KU55933 (10 μmol/L). Data are mean ± SEM.

### Chronic NNK exposure promotes persistent RAD51-BRCA2 activation but is associated with genomic instability

To determine the long-term consequences of NNK-induced DNA damage, we evaluated the activation of homologous recombination (HR) repair pathways following chronic NNK exposure. Immunofluorescence analysis revealed a marked decrease in nuclear RAD51 and BRCA2 foci in cells exposed to NNK for 50 days compared with control cells, indicating a lack of sustained activation of the HR DNA repair machinery in response to chronic genotoxic stress (Fig. 5A). Quantification of RAD51/BRCA2-positive cells over time demonstrated persistent accumulation of repair foci in NNK-treated cells, whereas control cells exhibited minimal HR activation (Fig. 5B). Despite the persistent presence of RAD51 and BRCA2 foci, chronic NNK exposure was associated with increased genomic instability, suggesting that prolonged genotoxic stress compromises the effectiveness of DNA repair. When considered together with the acute NNK response observed at 5 days, in which RAD51 and BRCA2 were efficiently recruited to DNA damage sites (Fig. 4), these findings indicate that chronic NNK exposure maintains DNA damage signaling but fails to fully preserve genomic integrity. We therefore propose a model in which acute NNK exposure activates ATM-γH2AX signaling and promotes RAD51-BRCA2-mediated DNA repair to maintain genomic stability. In contrast, chronic NNK exposure results in persistent DNA damage signaling despite continued RAD51 and BRCA2 accumulation, leading to impaired HR fidelity, unresolved DNA damage, and progressive genomic instability (Fig. 5C).

**Figure 5.**
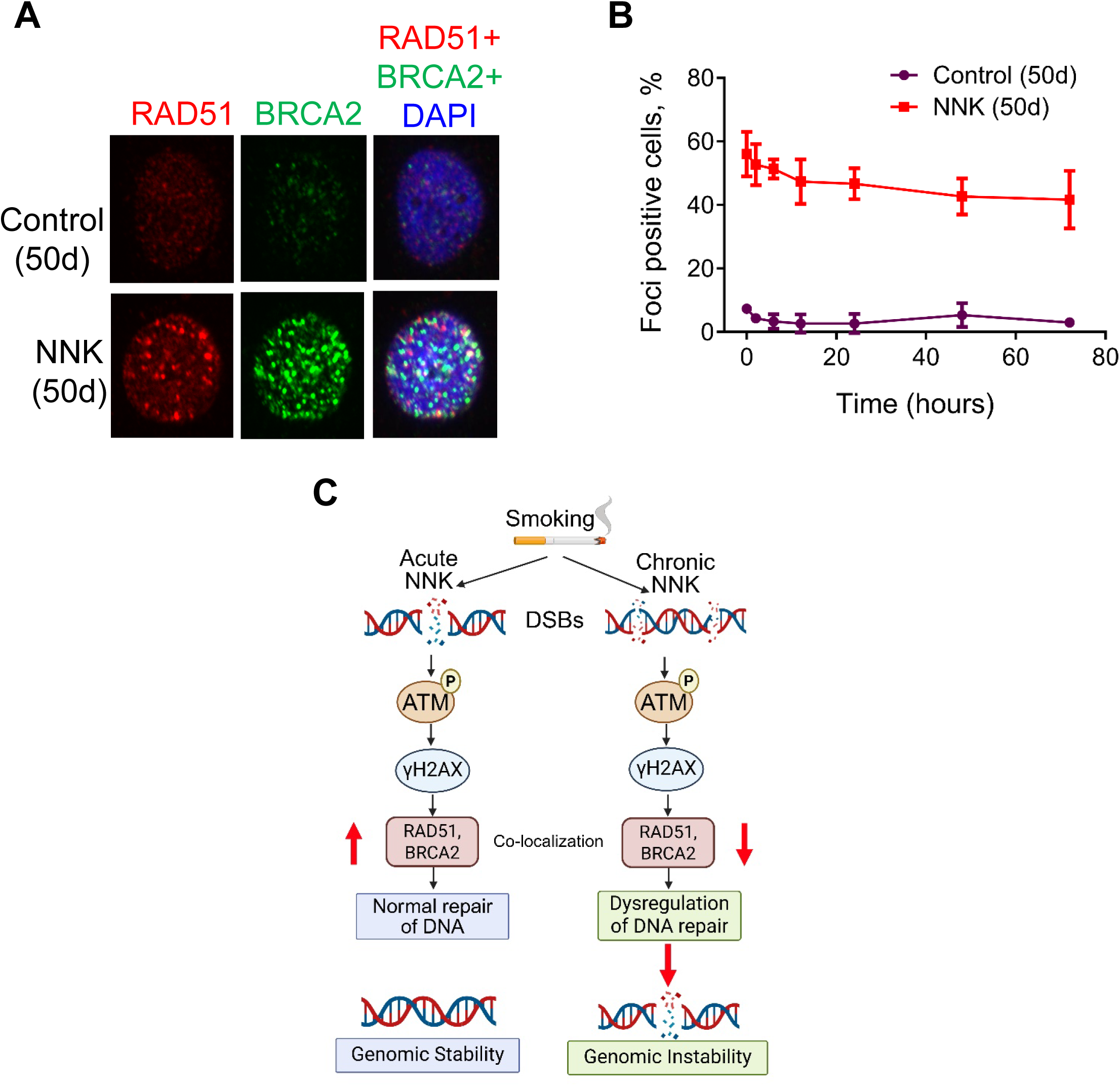
Chronic NNK induces chromosomal instability through homologous recombination repair deficiency. **(A**) Co-immunofluorescence of RAD51 and BRCA2 in H6c7 cells following 50-day chronic NNK treatment. Representative images are shown. (**B**) Persistence of RAD51/BRCA2 foci-positive cells (%) over 72 hours in control versus chronic (50-day) NNK-treated cells. Data are mean ± SEM. (**C**) Schematic summarizing how chronic NNK exposure promotes chromosomal instability through sequential DSB formation, ATM activation, HR gene upregulation, RAD51 cytoplasmic mislocalization, and defective HR repair.

### Elevated γH2AX expression is associated with higher tumor grade and shorter survival

To assess the clinical relevance of γH2AX expression in pancreatic cancer, we analyzed tissue microarrays (TMAs) containing normal pancreatic ducts, chronic pancreatitis, and pancreatic tumor specimens by immunohistochemistry. Representative images showed low γH2AX staining in normal pancreatic ducts, whereas pancreatic tumors exhibited markedly increased nuclear γH2AX expression, indicating enhanced DNA damage signaling in malignant tissues (Fig. 6A). Quantitative analysis of TMA specimens revealed a progressive increase in γH2AX expression across pancreatic disease states. Compared with normal ducts (n = 40), grade 2 (n = 37) and grade 3 (n = 10) tumors showed higher γH2AX staining (P = 0.012 and P = 0.0001, respectively), whereas chronic pancreatitis (n = 40) and grade 1 tumors (n = 21) did not differ significantly from normal ducts (P = 0.744 and P = 0.332). Among the 55 of 148 TMA patients with survival data, patients in the high γH2AX group (n = 33) had shorter overall survival than those in the low group (n = 22; median 11 vs. 36 months; log-rank P = 0.0007; Fig. 6C). Because this analysis was unadjusted and limited to the subset with survival data, it supports an association with overall survival but does not establish independent prognostic value. In exploratory TCGA-PAAD median-split analyses, higher RAD51 and BRCA2 mRNA expression was associated with shorter overall survival (P = 0.0018 and P = 0.018, respectively), whereas H2AX mRNA expression was not (P = 0.22; Supplementary Fig. 3A-C). Collectively, these findings support γH2AX as a marker of DNA damage signaling associated with higher tumor grade and adverse clinical outcome

**Figure 6.**
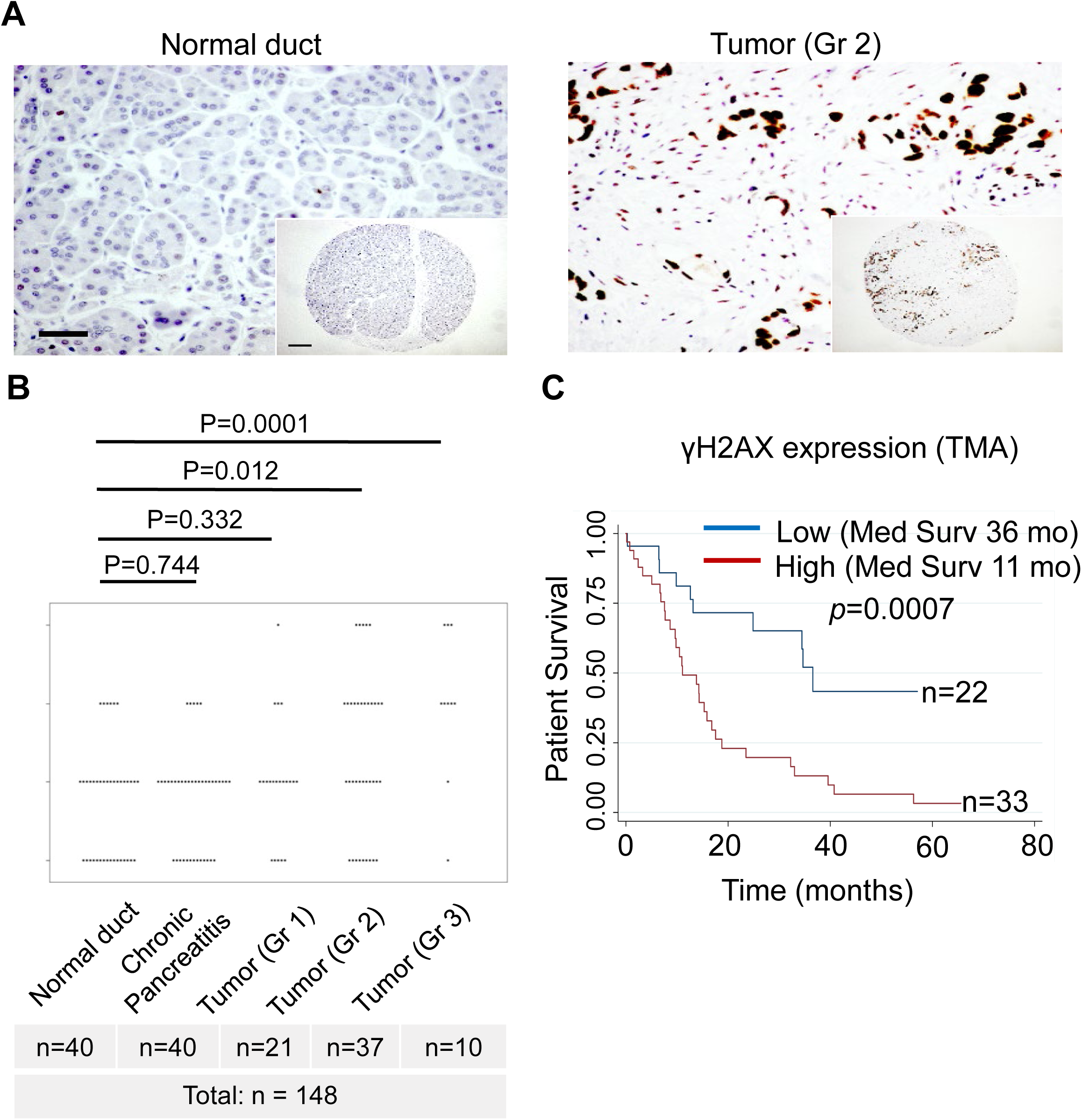
γH2AX protein expression correlates with tumor grade and predicts poor survival in PDAC. **(A**) Representative immunohistochemical images of γH2AX staining in normal pancreatic duct and PDAC (representative Grade 2 tumor shown). Human PDAC tissue microarray (TMA; n = 148 patients). Scale bar, 50μm. (**B**) Quantification of γH2AX staining index showing a progressive increase from normal ducts to Grade 1, Grade 2, and Grade 3 tumors. Statistical significance was assessed by Kruskal–Wallis test with Dunn’s post-hoc correction; pairwise p values are shown. Data are presented as dot plots with median and interquartile range. (**C**) Kaplan-Meier overall survival curves for PDAC patients stratified by γH2AX protein expression (high, n = 33; low, n = 22) assessed by IHC on the human PDAC TMA. High γH2AX protein expression is significantly associated with shorter overall survival (median 11 vs. 36 months; log-rank p = 0.0007). Note that this survival association is based on γH2AX protein quantification by IHC.

## DISCUSSION

Cigarette smoking is among the strongest environmental risk factors for PDAC and accounts for approximately 20–25% of pancreatic cancer cases worldwide (3, 4). Although tobacco carcinogens have long been implicated in pancreatic tumorigenesis, the molecular mechanisms linking smoking exposure to genomic instability and malignant transformation remain incompletely understood. In the present study, we demonstrate that the tobacco-specific nitrosamine NNK elicits a biphasic DNA damage response characterized by acute ATM-γH2AX activation and RAD51-BRCA2 recruitment, followed by persistent DDR signaling, altered RAD51 localization and chromosomal instability during chronic exposure. Through integrated analyses of transcriptomic datasets, mechanistic in vitro studies, genetically engineered mouse models, human pancreatic tissues, and TCGA datasets, our findings provide a comprehensive framework linking tobacco carcinogen exposure to pancreatic carcinogenesis and disease progression.

NNK undergoes metabolic activation via cytochrome P450-dependent pathways, generating reactive intermediates that form pyridyloxobutyl and methyl DNA adducts, including O^6^-methylguanine and O^4^-methylthymidine lesions (6). These lesions reduce replication fidelity and generate replication-associated DSBs. Consistent with this mechanism, we observed robust induction of DNA damage after NNK exposure, as shown by comet assays, elevated γH2AX levels, and co-localization of γH2AX with phosphorylated ATM. Pharmacologic inhibition of ATM using KU55933 significantly attenuated γH2AX accumulation, confirming ATM as a primary mediator of the NNK-induced DDR (11, 14). Our findings reinforce the utility of γH2AX as a functional biomarker of ongoing DNA damage and genomic stress within pancreatic epithelial cells.

A central finding of this study is the temporal dissociation between DNA damage induction and activation of the HR repair program. Acute NNK exposure generated substantial DSBs without significant transcriptional modulation of HR-associated genes, whereas chronic exposure elicited coordinated upregulation of multiple DNA repair components including RAD51, BRCA1, FANCD2, PALB2, XRCC1, CHEK2, PCNA, and PLK1. These observations suggest that persistent unresolved DNA damage is required to engage a broader adaptive DDR transcriptional response. Similar adaptive responses have been reported in cells exposed to chronic replication stress, where prolonged DDR activation enables survival under continuous genotoxic pressure while simultaneously facilitating accumulation of oncogenic alterations (10, 23, 24). Notably, the magnitude of RAD51 induction observed after prolonged NNK exposure is comparable to that reported in BRCA-deficient and replication stress-driven tumors (25, 26). Collectively, these findings suggest that chronic tobacco carcinogen exposure creates a state of persistent DDR activation that may contribute to early oncogenesis during pancreatic neoplastic evolution.

Despite robust transcriptional activation of the HR machinery, our data indicate that HR repair becomes functionally compromised following chronic NNK exposure. An intriguing finding was the altered subcellular localization of RAD51 following NNK exposure. Although total RAD51 expression increased substantially, a significant fraction accumulated in the cytoplasm rather than localizing to sites of nuclear DNA damage. Because RAD51 filament formation within the nucleus is essential for HR-mediated repair, loss of nuclear RAD51 localization is sufficient to confer functional HR deficiency despite elevated protein abundance (18, 19). This phenotype may arise from disruption of BRCA2-dependent RAD51 nuclear import and chromatin loading mechanisms (18, 27). Chromosomal instability and structural genomic rearrangements are defining features of PDAC and are strongly associated with disease progression, metastatic behavior, and therapeutic resistance (16, 28–30). Integrated genomic analyses have further demonstrated that alterations in DNA damage repair pathways contribute substantially to the genomic complexity and molecular heterogeneity of PDAC (28, 31). Together, these findings support a model in which chronic NNK exposure induces a functional “BRCAness-like” phenotype by impairing the RAD51-BRCA2 repair axis. The concept of BRCAness describes tumors that exhibit HR deficiency despite lacking canonical BRCA1 or BRCA2 mutations and has emerged as a critical driver of genomic instability in several malignancies (20, 25). Our findings extend this concept by demonstrating that chronic tobacco carcinogen exposure may generate a functional HR-deficient state through RAD51 mislocalization and defective repair complex assembly, thereby promoting genomic instability during pancreatic carcinogenesis.

Our analyses of human pancreatic tissues support the clinical relevance of persistent DDR signaling. Strong γH2AX staining was more frequent among smokers than non-smokers, and the association remained strong after stratification by tissue category using the counts shown in Fig. 3E. In the separate disease/grade analysis, grade 2 and grade 3 tumors had higher γH2AX staining than normal ducts. High γH2AX protein expression was also associated with shorter overall survival in the 55 patients with available survival data. This survival analysis was retrospective, univariate, and limited to a subset of the TMA cohort, so independent prognostic value remains to be established (13, 32). In contrast, H2AX mRNA expression was not associated with overall survival in TCGA-PAAD (Supplementary Fig. 3C), emphasizing that phosphorylated H2AX protein and H2AX transcript abundance are biologically distinct measurements The public genomic and transcriptomic analyses should also be interpreted as supportive rather than definitive. Supplementary Figure 2 includes two TCGA-PAAD representations derived from the same underlying source cohort; therefore, these entries are not independent and should not be pooled as separate cohorts to estimate mutation prevalence. In TCGA-PAAD, the median-split Kaplan-Meier analyses for RAD51, BRCA2, and H2AX are exploratory. Modeling expression continuously in Cox regression and adjusting for available clinical factors such as stage and age would provide a stronger assessment of independent association with outcome. These limitations do not affect the experimental observation of NNK-induced DDR signaling, but they constrain the strength of the clinical prognostic claims.

Our findings further suggest that DDR dysregulation in tobacco-associated PDAC extends beyond simple DNA damage accumulation and reflects broader alterations in genome maintenance programs. The coordinated upregulation of HR-associated genes, together with impaired RAD51 localization, highlights a disconnect between repair pathway activation and repair pathway competence. Such discordance may permit survival of genetically damaged cells while simultaneously accelerating the acquisition of oncogenic chromosomal alterations. Similar mechanisms have been implicated in the evolution of genomically unstable cancers, where chronic DDR signaling paradoxically promotes adaptation to replication stress and facilitates tumor progression (25, 26). The observation that RAD51 protein abundance increases despite defective nuclear localization underscores the importance of evaluating functional DNA repair capacity rather than relying solely on gene expression signatures.

Our in vivo studies further support the biological relevance of NNK-induced DNA damage during pancreatic tumorigenesis. We observed progressive accumulation of γH2AX throughout PanIN development and invasive PDAC formation in the KC mouse models. Importantly, NNK administration accelerated γH2AX accumulation, suggesting that tobacco carcinogens cooperate with oncogenic Kras signaling to amplify genomic stress during neoplastic progression. The concurrent increase in γH2AX in CK19-positive ductal epithelial cells and PanIN lesions confirms that DNA damage is associated with PDAC progression. These observations align with previous reports showing synergistic interactions between environmental carcinogens and mutant KRAS during pancreatic cancer development (8, 9, 33).

While our study provides comprehensive mechanistic and translational insights, several avenues remain for further investigation. Validation of the transcriptomic findings across additional pancreatic epithelial models, patient-derived organoids, and primary human tissues will broaden their applicability. The human tissue cohorts were retrospective and lacked comprehensive smoking exposure metrics such as pack-years. Prospective clinical studies incorporating detailed smoking exposure histories will further refine the relationship between tobacco exposure and DDR activation. Furthermore, although our data strongly support an association between chronic DDR activation and genomic instability, additional mechanistic studies are required to establish the precise molecular basis of RAD51 cytoplasmic sequestration and its contribution to HR dysfunction. Future investigations should focus on defining how chronic tobacco exposure alters nuclear transport pathways, protein-protein interactions, and DNA repair complex assembly during pancreatic tumor initiation and progression. In conclusion, this study provides a comprehensive mechanistic framework through which the tobacco carcinogen NNK promotes pancreatic carcinogenesis. We identify a temporally distinct DDR program characterized by ATM-γH2AX activation followed by chronic HR dysregulation, RAD51 mislocalization, and chromosomal instability. Across cellular, murine, human tissue, and PDAC patient cohorts, γH2AX emerges as a robust marker of DNA damage burden and adverse clinical outcome, while RAD51 mislocalization represents a previously underappreciated mechanism of functional HR deficiency induced by chronic tobacco carcinogen exposure. Collectively, these findings establish chronic tobacco-associated DDR dysregulation as a fundamental driver of pancreatic cancer progression and support γH2AX expression and RAD51 localization as clinically relevant biomarkers of genomic instability and DDR dysfunction in PDAC. Moreover, our data suggest that tobacco carcinogen exposure may represent a previously underrecognized mechanism for generating functional HR deficiency and BRCAness-like phenotypes in pancreatic cancer, thereby contributing to the molecular heterogeneity of PDAC described in recent genomic and transcriptomic studies (7, 26, 27, 34, 35).

## MATERIALS AND METHODS

### Cell Lines

Murine pancreatic intraepithelial neoplasia (PanIN) cell lines were derived from the *LSL-Kras^G12D/+;^ Pdx1^Cre/+^* genetically engineered mouse (GEM) models of PDAC (kindly provided by Dr. Andrew Lowy) and maintained as previously described (8). The immortalized human pancreatic ductal cell line HPDE6-E6E7 (H6c7) was kindly provided by Dr. M.S. Tsao (36, 37) and was maintained in keratinocyte growth medium (Invitrogen) supplemented with human epidermal growth factor and bovine pituitary extract.

### NNK Treatment Protocol

4-(Methylnitrosamino)-1-(3-pyridyl)-1-butanone (NNK; Toronto Research Chemicals, catalog no. N515000) was dissolved in dimethyl sulfoxide (DMSO; Sigma-Aldrich) to prepare a 100 mmol/L stock solution, aliquoted, and stored at −80°C protected from light. We prepared working concentrations fresh in complete growth medium immediately before each treatment. For acute exposure experiments, cells were treated with NNK at 0, 1, or 10 μmol/L for 5 days, refreshing the medium and NNK every 48 hours. For chronic exposure experiments, cells were treated continuously with NNK at 1 μmol/L for 50 days, with medium and NNK refreshed every 48 hours and cells passaged at approximately 80% confluence. DMSO vehicle (0.01% v/v final concentration) served as the negative control in all experiments. The 1 μmol/L concentration was selected based on published pharmacologically relevant concentrations in pancreatic tissue and prior studies from our laboratory (8, 38).

### Comet Assay

We performed the comet assay using the R&D Systems (#4250-050-K) (Trevigen kit) according to the manufacturer’s instructions. Briefly, we suspended cells in agarose, layered them onto slides (500 cells per slide), lysed them, and subjected them to electrophoresis under neutral conditions. We then fixed, dried, and stained samples with SYBR Gold. Images were acquired, and moments were calculated in ImageJ using a macro provided by Robert Bagnell (http://www.med.unc.edu/microscopy/resources/imagej-plugins-and-macros/comet-assay).

### Western blot Analysis and Antibodies

Cell lysates were collected in RIPA lysis buffer (50 mM Tris/HCl, pH 8.0, 250 mM NaCl, 1% NP40, 0.5% [w/v] sodium deoxycholate, 0.1% sodium dodecyl sulfate, complete mini protease and phosphatase inhibitors; the latter 2 were from Roche). Lysates were sonicated and centrifuged at 10,000 g for 15 minutes at 4^0^C. For each lane, we separated 35 µg of whole-cell lysate on NuPAGE Novex 4-12% Bis-Tris gels, transferred it onto an iBlot transfer stack and PVDF membranes, and used an iBlot dry blotting transfer system (Life Technologies). For immunodetection, we incubated membranes with the antibodies listed in Supplementary Table 1. We generated signals from HRP-coupled secondary antibodies by enhanced chemiluminescence (Millipore) and recorded them on Kodak film.

### γH2AX ELISA

The γH2AX ELISA assay was conducted to study the double-strand breaks using the γH2AX pharmacodynamics assay kit R&D Systems (#4418-096-K) (Trevigen kit), following the manufacturer’s instructions.

### Immunohistochemistry (IHC)

Formalin-fixed, paraffin-embedded (FFPE) tissue sections (4 μm) were deparaffinized in xylene and rehydrated through a graded ethanol series to distilled water. Antigen retrieval was performed by heating slides in 10 mmol/L sodium citrate buffer (pH 6.0) at 95°C for 20 minutes, followed by cooling to room temperature. Endogenous peroxidase activity was quenched with 3% hydrogen peroxide in methanol for 10 minutes. Sections were blocked with 5% normal goat serum in PBS containing 0.1% Tween-20 (PBS-T) for 1 hour at room temperature. Primary antibodies (see Supplementary Table S1) were diluted in 1% BSA/PBS-T and incubated overnight at 4°C. After three washes in PBS-T, sections were incubated with appropriate horseradish peroxidase (HRP)-conjugated secondary antibodies (1:500; Vector Laboratories) for 1 hour at room temperature. We developed the signal using the DAB Peroxidase Substrate Kit (Vector Laboratories) according to the manufacturer’s instructions. We counterstained sections with Mayer’s hematoxylin, dehydrated them, and mounted them with Permount (Fisher Scientific).

### Tissue microarray (TMA) of Pancreatic tissues

Human pancreatic tissue microarrays were designed and constructed as previously described (8, 38, 39). We conducted the studies in accordance with recognized ethical guidelines (Declaration of Helsinki, CIOMS, Belmont Report, U.S. Common Rule). γH2AX staining was scored by a board-certified pathologist (C. Shi) using a composite staining index calculated as the product of staining intensity score (0, no staining; 1+, weak; 2+, moderate; 3+, strong) and distribution score (0, no staining; 1+, <33% cells positive; 2+, between 33-66% cells positive; 3+, >66% cells positive), yielding a composite score of 0-9. For binary classification, we scored γH2AX as positive if any detectable nuclear staining was present. For survival analysis, we stratified patients by median staining index.

### Immunofluorescence (IF)

After the NNK treatment, cells were washed with PBS and fixed with 4% paraformaldehyde for 15 minutes at room temperature. These cells were washed 3 times with PBS and blocked/extracted for 30 minutes at room temperature in BlockAid blocking solution (Molecular Probes) and 0.3% Triton X-100 in PBS. Cells were incubated overnight at 4°C with primary antibodies diluted in 1% BlockAid blocking solution and 0.01% Tween-20 in PBS. Cells were washed 3 times for 15 minutes at room temperature with 0.01% Tween-20 in PBS (PBS-T). Appropriate secondary antibodies were conjugated for immunofluorescence with Alexa Fluor 488 and Alexa Fluor 568 for 1 hour of incubation at room temperature. Detailed information on primary antibodies and their dilutions is provided in Supplementary Table 1. The cells were washed 3 times in PBS-T and washed once in PBS and then washed in water and mounted with ProLong Gold plus DAPI (Invitrogen). All images were captured with an Olympus FV-1000 confocal microscope (Olympus), using a 60x oil immersion objective. The individual images were converted to tiff files with the FV-1000, and Photoshop (Adobe) was used to create the final figures. γH2AX foci or RAD51 foci were counted using Foci Counter (40).

### DSB Repair Assays and ATM Inhibition

Monoclonal γH2AX antibody detected with Alexa Fluor 594-conjugated goat anti-mouse secondary antibody (Molecular Probes, Eugene, OR) was utilized. For quantitative analysis, we counted foci in each cell using Foci Counter (40). In a single experiment, we usually counted cells until we counted at least 10 cells, and the number of foci varied. Antibody details are provided in Supplementary Table 1.

For ATM inhibition experiments, the selective ATM kinase inhibitor KU55933 (Selleckchem, catalog no. S1092) was dissolved in DMSO and used at a final concentration of 10 μmol/L. We added KU55933 to cells 1 hour before NNK treatment and maintained it throughout the treatment period. DMSO vehicle (0.01% v/v) served as the inhibitor control. γH2AX foci were quantified at 24 hours post-treatment as described above.

### Subcellular Fractionation

Nuclear and cytoplasmic protein fractions were prepared using the NE-PER Nuclear and Cytoplasmic Extraction Reagents kit (Thermo Fisher Scientific, catalog no. 78833) according to the manufacturer’s instructions. Briefly, cells were harvested by trypsinization, washed in ice-cold PBS, and lysed sequentially with CER I and CER II reagents to release the cytoplasmic fraction. Nuclear pellets were resuspended in NER reagent with intermittent vortexing. Protein concentrations of both fractions were determined by BCA assay. Equal protein amounts (20 μg) were resolved by SDS-PAGE and transferred to PVDF membranes (Millipore). Fractionation purity was confirmed by immunoblotting for Histone H3 (nuclear marker) and STAT3 (cytoplasmic marker). RAD51 distribution was assessed using anti-RAD51 antibody (Supplementary Table S1).

### RNA Extraction and Quantitative Real-Time PCR (qRT-PCR)

H6c7 cells were treated with DMSO (control) or NNK (1 µmol/L) for 50 days, and RNA was extracted using the Qiagen RNeasy kit (Qiagen) per manufacturer’s instructions. Vanderbilt Technology for Advanced Genomics (VANTAGE) assessed RNA quality. The qRT-PCR reactions analyzing BRCA1, BRCA2, FANCD2, PLK1, and RAD51 mRNA levels were performed using the Roche Transcriptor Universal cDNA master and analyzed on the LightCycler 480 II (Roche). Primers were ordered through Vanderbilt University DNA core. BRCA1: 5′-ttgttgatgtggaggagcaa-3′ and 5′-gattccaggtaaggggttcc-3′ with probe #22 (Roche); BRCA2: 5′-cctgatgcctgtacacctctt-3′ and 5′-gcaggccgagtactgttagc-3′ with probe #3 (Roche); FANCD2: 5′-cccagaactgatcaactctcct-3′ and 5′-ccatcatcacacggaagaaa-3′ with probe #29 (Roche); PLK1: 5′-tcctgaatgaagatctggaggt-3′ and 5′-aggagtcccacacagggtct-3′ with probe #7 (Roche); RAD51: 5′-tgagggtacctttaggccaga-3′ and 5′-cactgccagagagaccatacc-3′ with probe #7 (Roche); 18S: 5′-GCAATTATTCCCCATGAACG-3′ and 5-GGGACTTAATCAACGCAAGC-3′ (IDT) with probe #48 (Roche). Relative fold change of expression was calculated by the 2-ΔΔCt method.

### RNA Sequencing (RNA-Seq)

We processed H6c7 RNA-seq samples using standard methods on the Illumina HiSeq 2000 platform at the Vanderbilt Technologies for Advanced Genomics Next Generation Sequencing Core (http://vantage.vanderbilt.edu). Samples were sequenced in two multiplexed lanes of 100-bp single-end reads, yielding approximately 75 million mappable reads per lane. The Illumina pipeline was used for base calling and quality filtering. We performed subsequent analysis on the GALAXY open platform using the Tuxedo Tools analysis suite. We performed transcript assembly and abundance estimation (FPKM) with Cufflinks, and filtered transcripts with <0.5 FPKM. Differential gene and transcript expression, splice site use, transcription start site (TSS) use, and promoter use were assessed with Cufffdiff using a false discovery rate (FDR)-adjusted P-value (q-value), with q < 0.05 considered significant. Gene ontology analysis was performed with PANTHER, and Ingenuity analysis was conducted with assistance from Vanderbilt Technology for Advanced Genomics and Research Design, as previously described (8). Genome-wide ranked waterfall and fold-change distribution plots are shown in Supplementary Figure 1 A&B.

### Statistical Analysis

All statistical analyses were performed using GraphPad Prism (version 9.0; GraphPad Software, San Diego, CA) or R (version 4.3.0; R Foundation for Statistical Computing, Vienna, Austria) unless otherwise specified. All hypothesis tests were two-sided, with P < 0.05 considered statistically significant unless an FDR-adjusted q-value is reported. Data from independent biological experiments are summarized as mean ± SEM unless otherwise stated. Exact sample sizes and statistical tests are provided in the corresponding figure legends.

For two-group comparisons of continuous outcomes, we used two-sided unpaired Student’s t-tests; ordinal IHC staining indices were analyzed with the Mann-Whitney U test when a two-group comparison was required. For three or more independent groups, we used one-way ANOVA with Tukey’s post-hoc test for parametric outcomes and the Kruskal-Wallis test with Dunn’s correction for ordinal or non-parametric outcomes. Dose-response comparisons against the DMSO control used one-way ANOVA with Dunnett’s multiple-comparison test. Factorial experiments involving NNK and KU55933 were analyzed by two-way ANOVA, with multiplicity-adjusted pairwise comparisons where shown.

For the categorical smoking-status analysis shown in Fig. 3E, the association between smoking status and weak versus strong γH2AX staining was evaluated with a Cochran-Mantel-Haenszel test stratified by tissue category (normal duct, chronic pancreatitis, and tumor), and the common odds ratio with 95% confidence interval was reported. Differences in staining index across normal duct, chronic pancreatitis, grade 1, grade 2, and grade 3 groups using the Kruskal-Wallis test with Dunn’s post-hoc correction for multiple comparisons.

For the TMA overall-survival analysis, the 55 of 148 patients with available survival data were stratified into high and low γH2AX groups using the median staining index as the cutoff. Survival curves were compared by the log-rank (Mantel-Cox) test, and univariate Cox proportional hazards regression was used to estimate hazard ratios with 95% confidence intervals. Given the limited survival subset and absence of multivariable adjustment, this analysis was considered exploratory. TCGA-PAAD survival analyses were performed in R using the survival and survminer packages. For the Kaplan-Meier displays in Supplementary Fig. 3, patients were stratified at the median mRNA expression for each gene; these analyses were also considered exploratory.

RNA-seq differential expression analysis was performed with Cuffdiff (Cufflinks version 2.2.1) using an FDR threshold of q < 0.05. Fold changes are reported as linear fold changes relative to the corresponding DMSO control. Genome-wide ranked waterfall and fold-change distribution plots were generated in Python (version 3.10) using matplotlib and seaborn and are shown in Supplementary Fig. 1

## ACKNOWLEDGEMENTS

The authors thank Dr. Christine M. Eischen, PhD (Thomas Jefferson University, USA) and Prof. Ganesh Nagaraju (Indian Institute of Science, India) for assay development and project initiation, respectively. We also thank Dr. Xiaodian Sun and Dr. Yugang Ban for the analysis from the Biostatistics and Bioinformatics Shared Resource of the Sylvester Comprehensive Cancer Center at the University of Miami Miller School of Medicine.

This study was supported by the R21 CA209536 and R01 CA262526 grants from the National Cancer Institute (NCI) of the National Institutes of Health (NIH), the Stanley Glaser Foundation Research Award (UM SJG 2017-24), the James Esther and King Biomedical Research Program (22K06), and Florida Cancer Innovation Fund (25C22) of the Florida Department of Health, awarded to N.S. Nagathihalli. The Histopathology Core Service was conducted with the assistance of the Sylvester Comprehensive Cancer Center support grant, under the supervision of N. Nagathihalli. The research reported in this publication was supported by Sylvester Comprehensive Cancer Center and in part by the NCI of the NIH under Award Number P30 CA240139. The authors bear full responsibility for the content and the opinions expressed in this work, which may not necessarily reflect the official perspectives of the NIH.

## AUTHOR CONTRIBUTIONS

VK, VKG, SJ, SS, and NN conceived and designed the study, supervised all experiments, and wrote the manuscript. VK, VKG, SS, and KH performed NNK treatment experiments, comet assays, and γH2AX ELISA. VK, VKG, SS, and KH performed immunofluorescence, confocal microscopy, and foci quantification. VK, VKG, SS, and KH performed IHC on human TMA and mouse tissue sections. VK, VKG, SS, and KH performed RNA extraction, qRT-PCR, and RNA-seq library preparation. CS performed blind scoring of immunohistochemistry staining. PP performed RNA-seq bioinformatics analysis, TCGA-PAAD survival analysis, and cBioPortal data analysis. XC conducted statistical analysis; AD and NBM edited the manuscript. All authors reviewed and approved the final manuscript.

## ETHICS STATEMENT

All animal experimental protocols were approved by the Institutional Animal Care and Use Committee (IACUC) of the University of Miami and conducted in accordance with the Association for the Assessment and Accreditation of Laboratory Animal Care guidelines (Miami, FL, Protocol no. 18-081 and 21-093).

## DATA AVAILABILITY STATEMENT

We obtained TCGA-PAAD gene expression, somatic and germline alterations, and clinical data analyzed in this study from cBioPortal (https://www.cbioportal.org), and they are publicly available. All other data supporting the findings of this study are included in the article or available from the corresponding author upon reasonable request.

