## Supplementary figures and images for "Tobacco Carcinogen Induces Persistent DNA Damage and Homologous Recombination Dysregulation in Pancreatic Ductal Carcinogenesis"

### Supplementary figures

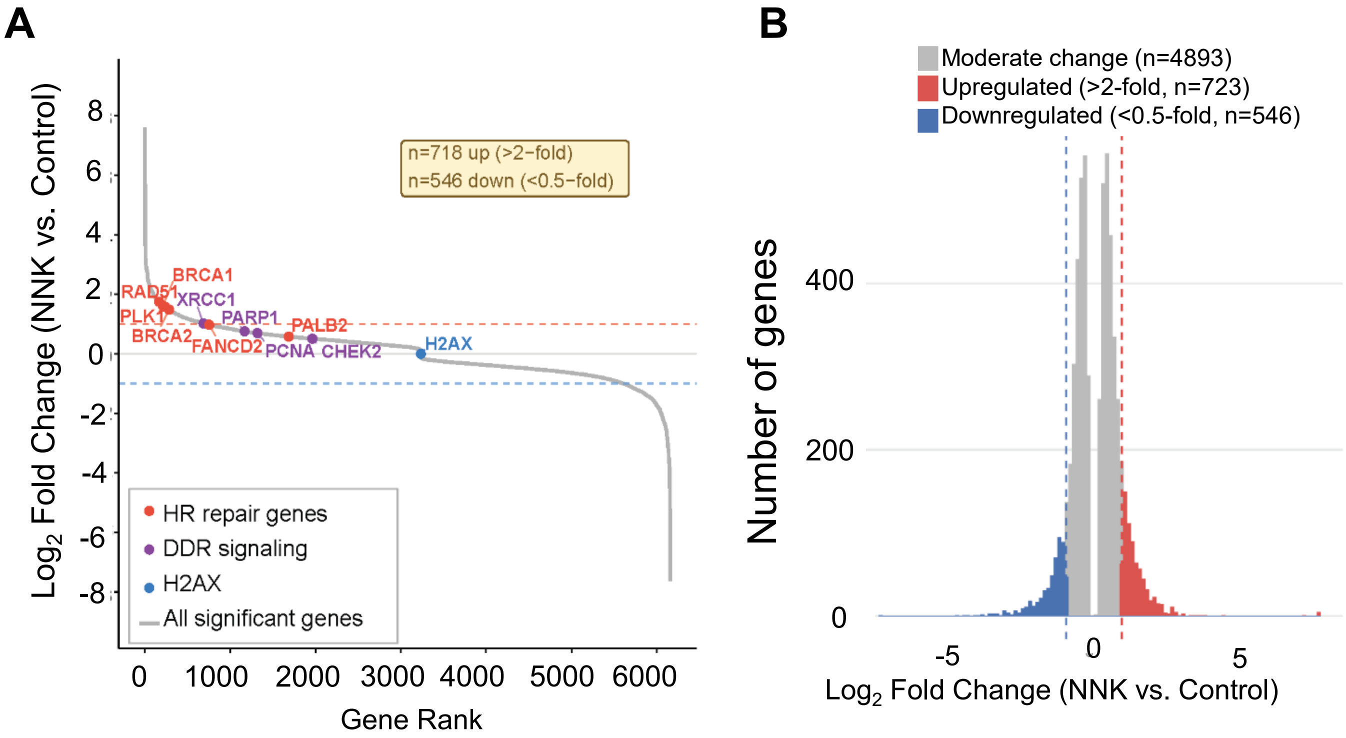


**Supplementary Figure 1**

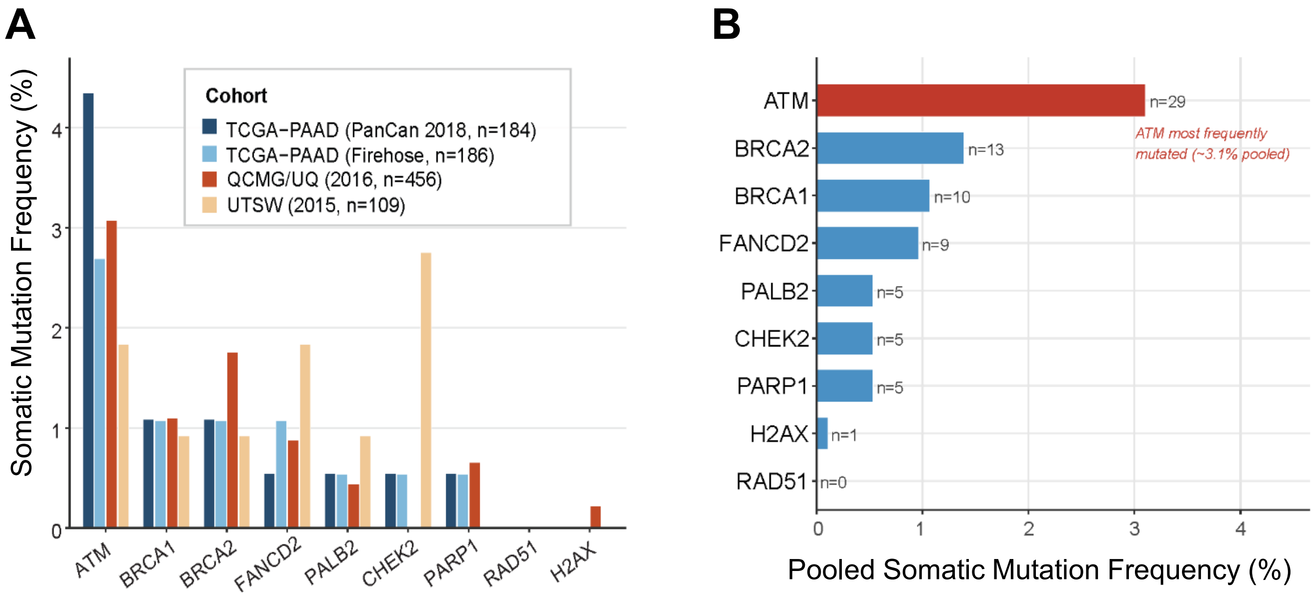


**Supplementary Figure 2**

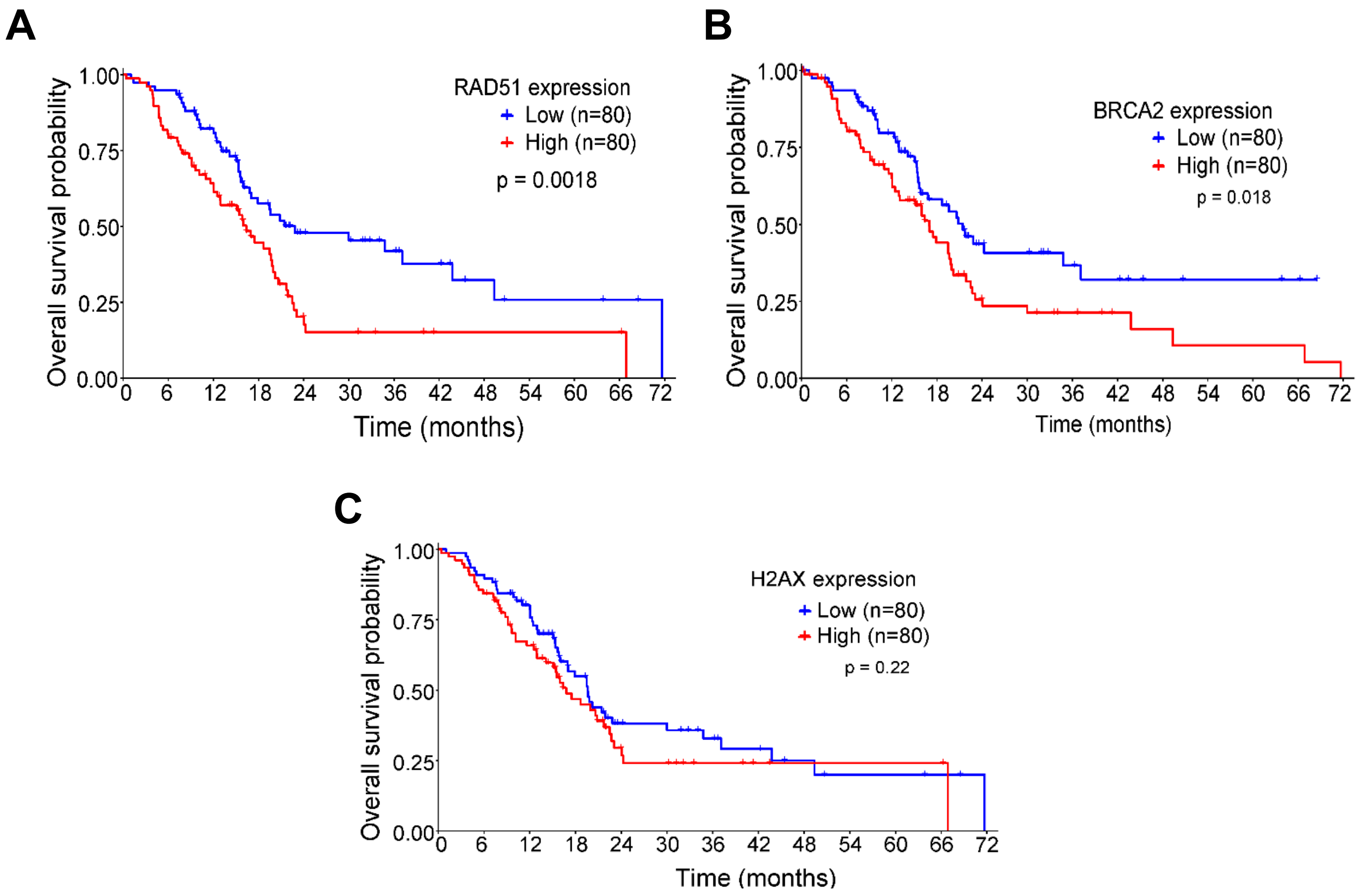


**Supplementary Figure 3**
