## Supplementary figure legends for "Tobacco Carcinogen Induces Persistent DNA Damage and Homologous Recombination Dysregulation in Pancreatic Ductal Carcinogenesis"

### Supplementary Figures

**Supplementary Figure 1. Genome-Wide Transcriptomic Alterations Induced by Chronic NNK Exposure**

**(A)** Ranked waterfall plot showing distribution of significant gene expression changes across the transcriptome following NNK treatment. **(B)** Distribution of log2-fold changes demonstrating widespread transcriptional reprogramming induced by chronic NNK exposure. We performed RNA-sequencing on independent biological replicates and conducted differential expression analysis using standard FDR-adjusted statistical methods.

**Supplementary Figure 2. Somatic Alterations in DNA Damage Response Genes Across Pancreatic Cancer Cohorts**

**(A)** Frequency of somatic mutations in major DDR genes across four independent PDAC cohorts. **(B)** Pooled analysis demonstrates the overall prevalence of DDR pathway alterations in pancreatic cancer. We obtained mutation frequencies from publicly available genomic datasets and analyzed them across all cohorts.

**Supplementary Figure 3. Elevated RAD51 and BRCA2 Expression Predicts Poor Clinical Outcome in PDAC**

Kaplan-Meier survival analysis based on RAD51 (**A**), BRCA2 (**B**), and H2AX (**C**) expression in patients from the TCGA-PAAD cohort. We stratified patients by gene expression levels and analyzed survival differences using the log-rank test.
