## Supplementary table for "Tobacco Carcinogen Induces Persistent DNA Damage and Homologous Recombination Dysregulation in Pancreatic Ductal Carcinogenesis"

**Supplementary Table 1. Antibodies used in this study.**

Primary antibodies used for immunohistochemistry (IHC), immunofluorescence (IF), and western blotting (WB) are listed below. Catalog numbers, dilutions, and applications are provided.

| **Target** | **Clone/Cat. No.** | **Host** | **Company** | **Dilution (IHC)** | **Dilution (IF)** | **Dilution (WB)** |
|---|---|---|---|---|---|---|
| γH2AX (pSer139) | EP854(2)Y / ab81299 | Rabbit | Abcam | 1:200 | 1:200 | 1:1000 |
| H2AX (total) | 7631 | Rabbit | Cell Signaling Technology | — | — | 1:1000 |
| pATM (pSer1981) | 10H11.E12 / 4526 | Mouse | Cell Signaling Technology | — | 1:200 | 1:1000 |
| ATM (total) | D2E2 / 2873 | Rabbit | Cell Signaling Technology | — | — | 1:1000 |
| pCHK2 (pThr68) | C13C1 / 2197 | Rabbit | Cell Signaling Technology | — | — | 1:1000 |
| RAD51 | H-92 / sc-8349 | Rabbit | Santa Cruz Biotechnology | — | 1:100 | 1:500 |
| BRCA2 | Ab-1 / OP95 | Mouse | Millipore/ Calbiochem | — | — | 1:500 |
| CK19 | A53-B/A2 / M0888 | Mouse | Millipore/Dako | 1:100 | — | — |
| Histone H3 | D1H2 / 4499 | Rabbit | Cell Signaling Technology | — | — | 1:1000 |
| STAT3 | 124H6 / 9139 | Mouse | Cell Signaling Technology | — | — | 1:1000 |
